# Large Serine Integrase Off-Target Discovery with Deep Learning for Genome Wide Prediction

**DOI:** 10.1101/2024.10.10.617699

**Authors:** Matthew H. Bakalar, Thomas Biondi, Xiaoyu Liang, Didac Santesmasses, Anne M. Bara, Japan B. Mehta, Jie Wang, Dane Z. Hazelbaker, Jonathan D. Finn, Daniel J. O’Connell

## Abstract

Large Serine Integrases (LSIs) hold significant therapeutic promise due to their ability to efficiently incorporate gene-sized DNA into the human genome, offering a method to integrate healthy genes in patients with monogenic disorders or to insert gene circuits for the development of advanced cell therapies. To advance the application of LSIs for human therapeutic applications, new technologies and analytical methods for predicting and characterizing off-target recombination by LSIs are required. It is not experimentally tractable to validate off-target editing at all potential off-target sites in therapeutically relevant cell types because of sample limitations and genetic variation in the human population. To address this gap, we constructed a deep learning model named IntQuery that can predict LSI activity genome-wide. For Bxb1 integrase, IntQuery was trained on quantitative off-target data from 410,776 cryptic *attB* sequences discovered by Cryptic-seq, an unbiased in vitro discovery technology for LSI off-target recombination. We show that IntQuery can accurately predict in vitro LSI activity, providing a tool for *in silico* off-target prediction of large serine integrases to advance therapeutic applications.

## Introduction

The large serine integrase (LSI) family constitutes a diverse group of site-specific recombinases that play pivotal roles in mediating DNA rearrangements^1-3^. Serine integrases, in contrast to their tyrosine recombinase counterparts, utilize a serine residue for catalysis, leading to distinct mechanistic features^4^. This large family encompasses integrases with varying sizes and functionalities, with notable members including PhiC31 integrase from *Streptomyces* bacteriophage PhiC31^5^ and Bxb1 integrase discovered in mycobacteriophage Bxb1^6^. Both PhiC31 and Bxb1 integrases are well-recognized for their utility in site-specific recombination applications by virtue of direct recombination between phage attachment site *attP* and bacterial attachment site *attB* with the requirement of no co-factors or DNA supercoiling^7-9^. The precise and efficient DNA manipulation capabilities of the large serine integrase family have positioned it as an attractive tool for synthetic biology and genome editing applications^10-14^.

Large serine integrases facilitate recombination between attachment sites on linear or circular DNA substrates^5, 15^. Recently CRISPR-directed integrase editing strategies have been described^11, 14^ that enable programmable genomic insertion of DNA cargo to facilitate gene replacement strategies^16^. Unlike CRISPR/Cas9 knock-in approaches, which rely on the host cell’s DNA repair mechanisms^17-19^, LSI integration operates independently of host cell factors and does not require double-stranded breaks (DSBs). Instead, it directly integrates the template DNA, resulting in significantly higher fidelity compared to other error-prone gene writing methods^5, 15^. This approach to gene insertion has the advantage of minimizing unintended editing at the on-target locus, but it still presents the risk of potential off-target insertion and gross chromosomal rearrangements related to LSI-mediated recombination at ‘cryptic’ or ‘pseudo’ attachment sequences that may be present in the human genome^20-22^.

The rapid development of new medicines driven by the expansion of genome editing technologies^23^ has resulted in approved cures for sickle cell disease using ex vivo products^24, 25^. Additionally, promising clinical data has emerged for *in vivo* genome editing, where lipid nanoparticles encapsulating Cas9 mRNA and guide RNA are administered systemically^26, 27^. In response to these exciting advances in genetic medicines, the FDA has released non-binding recommendations for the assessment of safety, including potential genotoxicity from off-target editing events^28^. Nonclinical safety studies designed to discover potential risks should use multiple methods (e.g., *in silico*, biochemical and cellular-based assays) that include a genome-wide analysis to reduce bias in identification of potential off-target sites^28^.

To help safely advance CRISPR-directed integrases for clinical trials, we previously described two empirical genome-wide discovery technologies, HIDE-seq and Cryptic-seq^20^, to enable unbiased off-target discovery in isolated human genomic DNA (gDNA) samples. However, an effective rank-ordering *in silico* prediction tool for LSI off-target recombination has not been described. To address this gap, we constructed a deep learning model named IntQuery, using the empirical off-target discovery data from Cryptic-seq for the LSI Bxb1 integrase across multiple central dinucleotide targets. The quantitative nature of Cryptic-seq data from 410,776 potential off-target sites in the human genome allowed us to leverage a simple deep learning model to effectively predict and rank LSI off-target recombination potential genome-wide.

## Results

To determine if new methods for the computational prediction of potential off-target sites for an LSI were required to advance therapeutic CRISPR-directed integrases, we first evaluated HOMER^29^, a well-established DNA-protein interaction model based on position weight matrices (PWM). Utilizing the position weight matrix identified by HIDE-Seq^20^, HOMER’s default parameters predicted 4,598,283 potential off-target sites within the human genome (**Figure 1A**). This number of potential off-target sites is too high for effective verification of potential off-target editing in cells with the current state of technology. Therefore, we sought to leverage the wealth of empirical and quantitative off-target discovery data from the highly sensitive Cryptic-seq technology^20^ to develop IntQuery, a deep learning model for quantitative prediction of LSI off-target integration at any DNA sequence.

**Figure 1.**
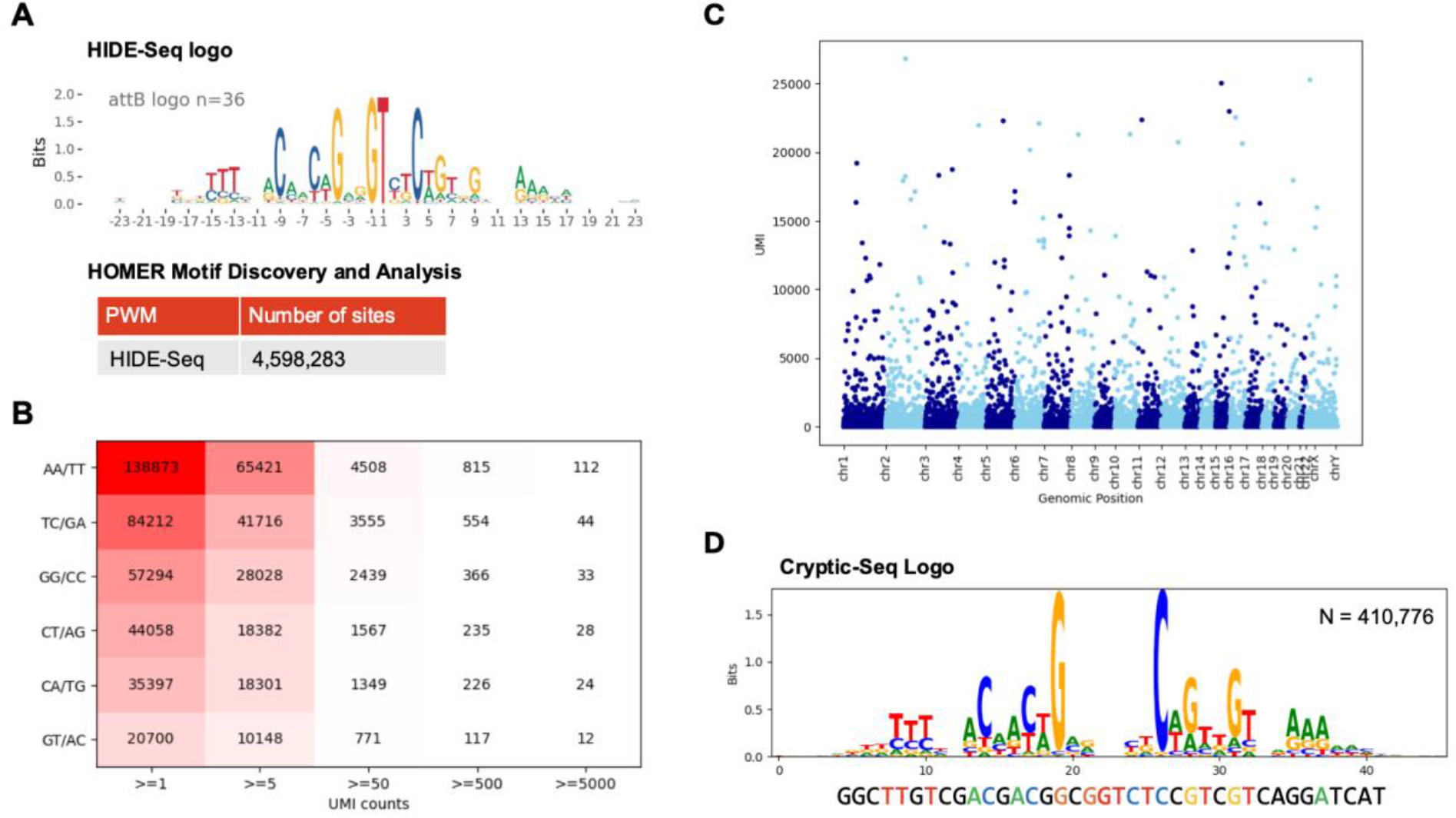
Cryptic-seq off-target discovery for the LSI Bxb1 across multiple central dinucleotides. **(A)** Genome-wide search with the PWM generated by HIDE-seq with HOMER identified 4,598,283 potential off-target loci. **(B)** Data matrix created with UMI count bins from >=1 UMIs up to >= 5000 UMIs of the number of cryptic *attB* sites detected in the human genome with DNA donor substrates containing an *attP* containing either AA, TC, GG, CT, CA, and GT central dinucleotides. **(C)** Genomic distribution of cryptic *attB* sites plotted against the UMI signal discovered by Cryptic-seq. **(D)** DNA sequence motif from the 410,776 cyptic *attB* sites discovered by Cryptic-seq. Natural Bxb1 *attB* sequence is displayed on the bottom.

LSI attachment sequences are demarcated by a canonical central dinucleotide^2^ which promotes annealing and ligation after 180-degree rotation to complete the recombination reaction. While the central dinucleotide does not directly interact with the LSI protein, it plays a critical role in determining recombination specificity because the landscape of cryptic attachment sites aligns with the central dinucleotide of the complementary attachment sequence. When applied to CRISPR-directed integrases, this feature allows simultaneous and specific multiplex gene insertion of unique cargos by utilizing orthogonal central dinucleotides^14^. Although there are 16 possible central dinucleotide combinations, 4 of the combinations are palindromic and permit bidirectional insertion, and 6 of the combinations represent complementary counter parts that allow directional control over the orientation of insertion. Therefore, we performed Cryptic-seq with substrates spanning the 12 central dinucleotides with directional insertion (AA/TT, TC/GA, GG/CC, CT/AG, CA/TG, GT/AC) to determine the off-target editing landscape of Bxb1. Cryptic-seq discovered a total of 410,776 unique potential off-target sites (**Figure 1B and Supplementary Table 1**) distributed throughout the human genome (**Figure 1C**). The DNA motif created from these discovery data was similar to the motif discovered by HIDE-seq^20^ and revealed the expected features of sequence conservation and palindromicity related to the dimerization of Bxb1 on attachment sequences (**Figure 1D**).

Cryptic-seq is a quantitative biochemical off-target discovery assay because each recombination event imparts a unique molecular identifier (UMI) to the NGS reads^20^. We reasoned that our empirical database of 410,776 cryptic attachment sites in the human genome could be used to train a machine learning model to predict integrase activity from a DNA sequence alone. To demonstrate this approach, we trained a simple multi-layer perceptron model with two hidden layers to perform a regression task, which we call IntQuery^30^. We trained IntQuery to predict log-transformed UMIs from a one-hot encoding of the cryptic attachment site sequence, replacing the central dinucleotide of each sequence with NN. To prevent oversampling low UMI sites (17% of sites have UMI = 1), we used UMI-weighted sampling of the training data for each epoch. We conducted five-fold cross-validation on the training data, achieving a Spearman correlation of ρ = 0.42 between the target values and predictions (**Figure 2A)**. A linear regression model trained with an identical protocol achieved a Spearman correlation of ρ = 0.34 across all predictions. Plotting target vs predicted values for both models revealed that while the MLP model maintains correlation across the entire range of UMI values, the linear regression model is unable to discriminate between low UMI and high UMI sites (MLP ρ = .51, linear regression ρ = 0.15 for sites with target values > 4). Therefore, IntQuery provides a simple method for predicting LSI cryptic attachment site identity and activity directly from DNA sequence. We anticipate that IntQuery will be valuable as an *in-silico* method for discovery of off-target sites and prioritization of potential high-activity sites for verification in LSI-edited cells of interest.

**Figure 2.**
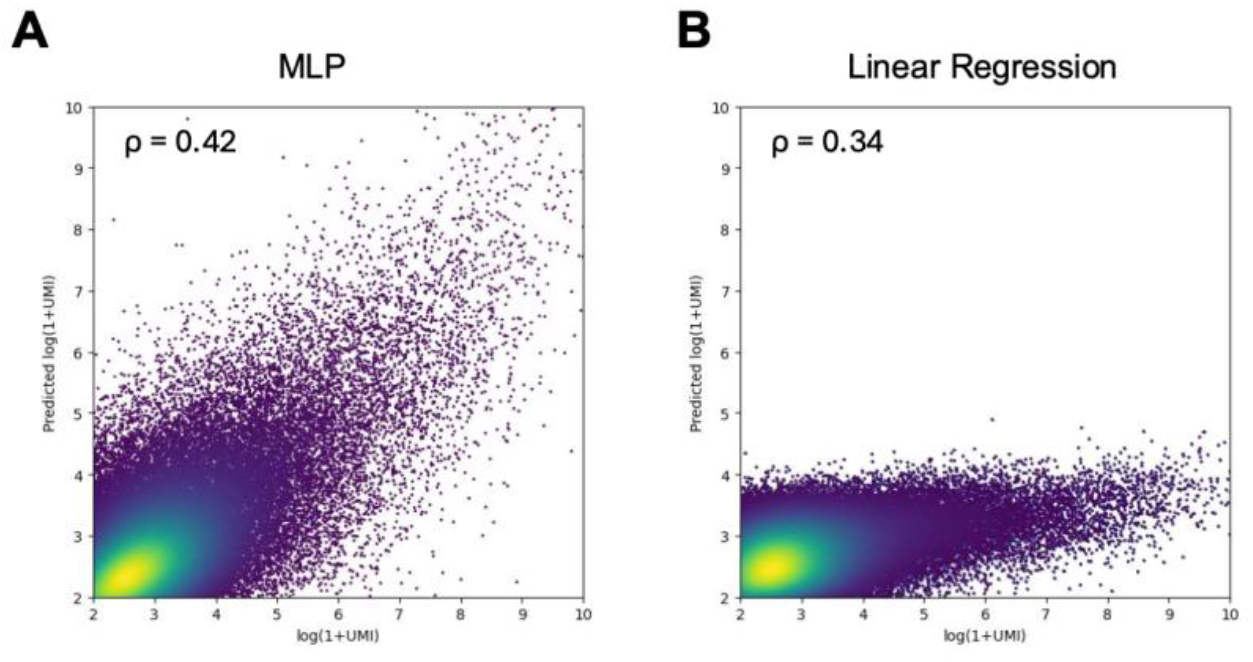
Target vs predicted cross-validation plots from IntQuery and a linear regression model. **(A)** Five-fold cross-validation between the target values and predictions of IntQuery revealed a Spearman correlation of ρ = 0.42. **(B)** A linear regression model trained with an identical protocol achieved a Spearman correlation of ρ = 0.34 across all predictions.

## Discussion

IntQuery is a machine learning approach to predict the potential for LSI recombination at any site in the human genome. The approach taken by IntQuery is applicable to empirical LSI off-target discovery data from any genome. Computational off-target prediction with IntQuery complements empirical discovery data generated in the laboratory to help scientists prioritize potential off-target sites for experimental verification in edited cells. This two-step strategy was inspired by the approaches pioneered for the first Cas9 genome editing therapies^24, 26, 31^, because the risk of off-target editing from Cas9 and LSIs are both dependent on DNA sequence homology^22, 32^.

IntQuery can also be used for variant-aware off-target prediction^33^ for any large serine integrase, including naturally discovered^12, 14^ and engineered ^21, 34, 35^ versions, by generating a large quantitative discovery dataset with technologies like Cryptic-Seq^20^. Experimental verification of IntQuery off-target predictions in edited cells will increase its relevance in supporting the non-clinical studies required for a new drug application. We hope our communication of a deep learning off-target prediction tool will help enable the safe development of LSI-based therapeutics.

## Methods

### Position Weight Matrix Prediction of Bxb1 Cryptic Sites with HOMER

The position weight matrix discovered by HIDE-Seq^20^ was used to predict locations in the hg38 human reference genome^36^ that might be loci for off-target recombination by Bxb1. Briefly, we aligned the sequences flanking the integration sites discovered by HIDE-seq^20^ and generated a custom motif file based on the position frequency matrix and ran HOMER v4.11 motif analysis (scanMotifGenomeWide.pl)^29^ with default parameters to predict binding at all sites in the hg38 human reference genome.

### Cryptic-seq

Cryptic-seq was performed as previously described^20^ with the following modifications. HEK293*attB*^20^ gDNA sheared in a ME220 focused-ultrasonicator (Covaris) to an average fragment length of approximately 350-500 bp. End prep and dA tailing of the gDNA was performed by using NEBNext® Ultra™ II DNA Library Prep Kit for Illumina® (E7645L) as protocol instructed, after which Illumina TruSeq sequencing annealed Y adaptors containing 8 bp UMIs were ligated to gDNA listed below.

Two independent cryptic-seq reactions with the following pools of cryptic-seq donor plasmids were performed. Reaction 1 contains the following plasmids at equimolar ratios: PL2327_AA, PL2341_TT, PL2337_GG, PL2332_CC, PL2339_TC, PL2335_GA, with a final total plasmid concentration of 30 nM. The plasmid pool was then reacted with 1 μg sheared/adapter-ligated gDNA and 1 μM Bxb1 integrase at 37°C for 4h in a recombination buffer^7^. Reaction 2 contains the following plasmids at equimolar ratios: PL2312_GT, PL2328_AC, PL2331_CA, PL2340_TG, PL2334_CT, PL2329_AG and was reacted under the same conditions as reaction described above. A representative sequence map of the cryptic-seq plasmid PL2312_GT can be found in our previous publication^20^ with all plasmids in this study being identical to PL2312 except for the indicated central attP dinucleotide. The reaction was stopped by adding sodium dodecyl sulfate to a final concentration of 0.1% and the products were cleaned using the Zymo clean and concentrator kit (Zymo research). PCR was used to amplify the regions of integration using Q5 polymerase (NEB) and common primer for P5 (LM_P5) and N7 primer (GN037_CrypticSeq_N7_i7_N701). Following PCR, a 1.5x AMPure bead clean-up was performed, and products were run on a Tapestation 4200 (Agilent Technologies) using D1000 tape to confirm amplification. Library concentration was determined using the Next Library Quant Kit (NEB) for Illumina. Libraries were sequenced on an Illumina NovaSeq (Fulgent Genetics).

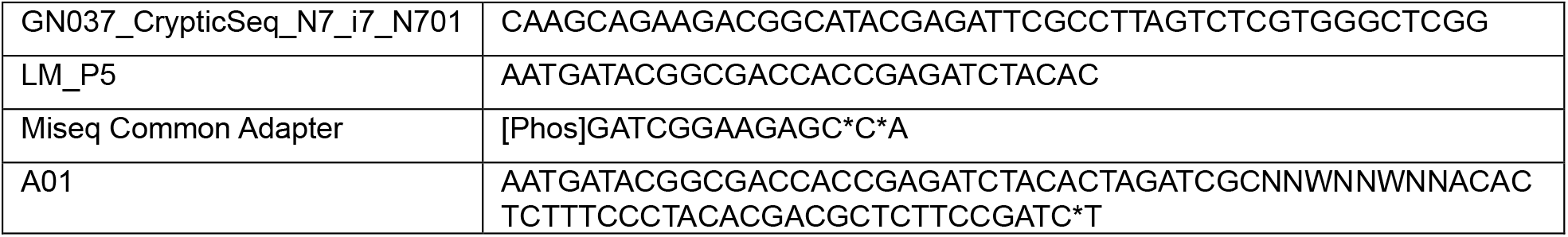

For bioinformatic analysis of Cryptic-seq data, FASTQ files from Illumina sequencing were loaded into a custom Cryptic-seq bioinformatic pipeline tbChaSIn developed by Tome Biosciences and Fulcrum Genomics to discover and quantify cryptic recombination sites from Cryptic-seq data (https://github.com/didacs/tbChaSIn). The bioinformatic workflow starts with the trimming of reads that contain leading *attP* or *attB* sequences (P, P’ or B, B’) at the 5’ end of R2. Untrimmed reads were discarded. Included reads were further trimmed to remove Illumina TrueSeq adapter sequence (AGATCGGAAGAGCGTCGTGTAGGGAAAGAGTGT) from the 3’ end of R2. Trimmed read sequences were aligned against the hg38 human reference genome^36^ with appended *attP* and *attB* sequences by BWA aligner^37^ to create mapped BAM files for each sample. The resulting BAM files were deduplicated by Picard^38^. The integration sites are identified by the first base of R2 and quantified by the number of deduplicated reads that pass mapping quality threshold (MAPQ > 20). Output .csv files containing sites per sample along with collation of sites across samples were generated.

### IntQuery

Cryptic attachment sequences were one-hot encoded and used as input to predict their corresponding UMI values. UMI values were scaled to log (1 + UMI) before training. The MLP model was written in Pytorch and consisted of an input layer, two hidden layers with dropout, and an output layer with a single dimension, with mean-squared-error between prediction and target used as an error function^39^. To prevent oversampling of low UMI sites, we randomly sampled the training data with a weight directly proportional to each sites log (UMI) value. For demonstration purposes in this paper, we trained a model with hidden dimensions of 200 and 5% dropout. In preliminary experiments we found that model performance is relatively insensitive to model architecture but don’t rule out the possibility that alternative models might outperform the simple MLP described here.

## Data and model availability

All code and training data have been made publicly available on GitHub: https://github.com/mhbakalar/intquery. The Cryptic-seq next-generation sequencing data is available under NCBI SRA bioproject PRJNA1169517. The code used for Bxb1 Cryptic-seq is available on GitHub https://github.com/didacs/tbChaSIn and the genomic coordinates (hg38) for discovered sites are detailed in **Supplementary Table1**.

## Acknowledgements

We would like to thank all members of Tome Biosciences for their support in the generation of this work.

